# The Evolution of Placental Invasion and Cancer Metastasis are Causally Linked

**DOI:** 10.1101/528646

**Authors:** Kshitiz, Junaid Afzal, Jamie D. Maziarz, Archer Hamidzadeh, Cong Liang, Eric M. Erkenbrack, Hong Nam, Jan-Dirk Haeger, Christiane Pfarrer, Thomas Hoang, Troy Ott, Thomas Spencer, Mihaela Pavlicev, Doug Antczak, Andre Levchenko, Günter P. Wagner

**Affiliations:** Yale Institute of Systems Biology, Yale University, West Haven, CT; Department of Biomedical Engineering, Yale University, New Haven, CT; Department of Biomedical Engineering, University of Connecticut Health Center, Farmington, CT; Department of Ecology and Evolutionary Biology, Yale University, New Haven, CT; Department of Medicine, University of California San Francisco, CA; Center for BioMicrosystems, Korea Institute of Science and Technology, S. Korea; Institute of Anatomy, University of Veterinary Medicine, Hannover, Germany; Department of Animal Science, Center for Reproductive Biology and Health, Penn State University, University Park, PA.; Division of Animal Sciences, University of Missouri, Columbia, MO; Cincinnati Children’s Hospital and Medical Center, Cincinnati, OH; Department of Microbiology and Immunology, Cornell University, Ithaca, NY; Department of Obstetrics, Gynecology and Reproductive Sciences, Yale Medical School, New Haven, CT; Department of Obstetrics and Gynecology, Wayne State University, Detroit, MI

## Abstract

Among mammals, the extent of placental invasion is correlated with vulnerability to malignancy. Animals with more invasive placentation (e.g. humans) are more vulnerable to malignancy, whereas animals with a non-invasive placenta (e.g. ruminants) are less likely to develop malignant cancer. To explain this correlation, we propose the hypothesis of *Evolved Levels of Invasibility* (ELI) positing that the permissiveness of stromal tissue to invasion is a unitary character affecting both placental and cancer invasion. We provide evidence for this hypothesis by contrasting invasion of human and bovine cancer and placental cells into a lawn of stromal cells from different species. We find that both bovine endometrial and skin fibroblasts are more resistant to invasion of placental and cancer cells than their human counterparts. Gene expression profiling identified genes with high expression in human but not bovine fibroblasts. Knocking down of a subset of them in human fibroblasts leads to significantly stronger resistance to cancer cell invasion. Comparative analysis of gene expression among mammals suggests that humans evolved higher vulnerability to malignancy than the eutherian ancestor, possibly as a correlate of more invasive placentation, and boroeutherians evolved to decrease stromal invasibility. Identifying the evolutionary determinants of stromal invasibility can provide significant insights to develop rational anti-metastatic therapeutics.

## Introduction

Placental invasion into the maternal endometrium of the uterus exhibits substantial similarities to early cancer dissemination into stroma^1-4^. The stromal compartment surrounding the tumor is considered a natural barrier, which cells disseminating from a neoplastic lesion need to breach to become malignant. These similarities are both physiological and molecular, and have inspired the hypothesis of antagonistic pleiotropy^5,6^. According to this hypothesis, trophoblasts acquired the capacity to invade into the endometrial tissue leading to the development of invasive placentation, i.e. in the stem lineage of placental (eutherian) mammals. Consequently, these mechanisms became activated in cancer cells, leading to metastasis. This hypothesis implies that cancer malignancy evolved coincidental with, or after invasive placentation, and thus should be limited to placental mammals. This prediction, however, is inconsistent with various observations ^7^. For instance, marsupial mammals, like opossums, get invasive skin cancers, even though marsupials do not belong to the placental mammal clade and their placenta is ancestrally non-invasive^8,9^. Here we pursue an alternative scenario in which stromal cells of the uterus evolved to either resist or permit invasion, rather than a model being driven by the evolution of placental invasiveness.

Further evidence that evolution of cancer malignancy is a stromal cell phenomenon comes from the fact that the molecular mechanisms employed by cancer cells to metastasize are shared with other biological processes, e.g., those regulating gastrulation, wound healing, extravasation by leukocytes etc., and are not exclusive to trophoblast invasion into the endometrium^6,10,11^. Invading cancer cells employ mechanisms that evolved much earlier than placental invasion, and therefore, the evolution of invasive placentation per se cannot be responsible for the origin of malignant cancer. It is important to note, however, that the invasiveness of the placenta continued to evolve after the origin of placental development. Placental invasion reverted to a non-invasive phenotype in many lineages of placental mammals, as well as evolved into an even higher degree of invasiveness in the great apes, which includes humans^12-14^. A complete loss of placental invasion evolved in the hoofed mammals, like cows and horses and their relatives, and it is exactly in those animals that lower malignancy rates for a variety of cancers have been reported^7^. In contrast, humans with very invasive placentas are highly vulnerable to melanoma malignancy.

Based on the evidence outlined above, here we here advance the hypothesis that evolution in the permissiveness, or resistance of stromal cells to placental invasion is mechanistically linked to the vulnerability to cancer malignancy. We term this hypothesis *Evolved Levels of Invasibility (*ELI) (Figure 1A). We experimentally test the ELI hypothesis by engineering the interface between the invasive epithelial cells facing stromal cells in an environment mimicking a structured extracellular matrix (ECM) (Figure 1B). Using this experimental model, we demonstrate that human fibroblasts are more permissive to invasion by placental trophoblast as well as cancer cells compared to their bovine counterparts, reproducing the *in vivo* differences. We then identify some of the factors responsible for the evolved resistance to invasion in bovine fibroblasts, presenting a genetic roadmap to target metastasis by increasing stromal resistance to invasion. This study also highlights how investigating evolutionarily driven mechanisms may lead to the identification of therapeutic targets, pointing to the clinical potential of evolutionary comparative analysis.

**Figure 1.**
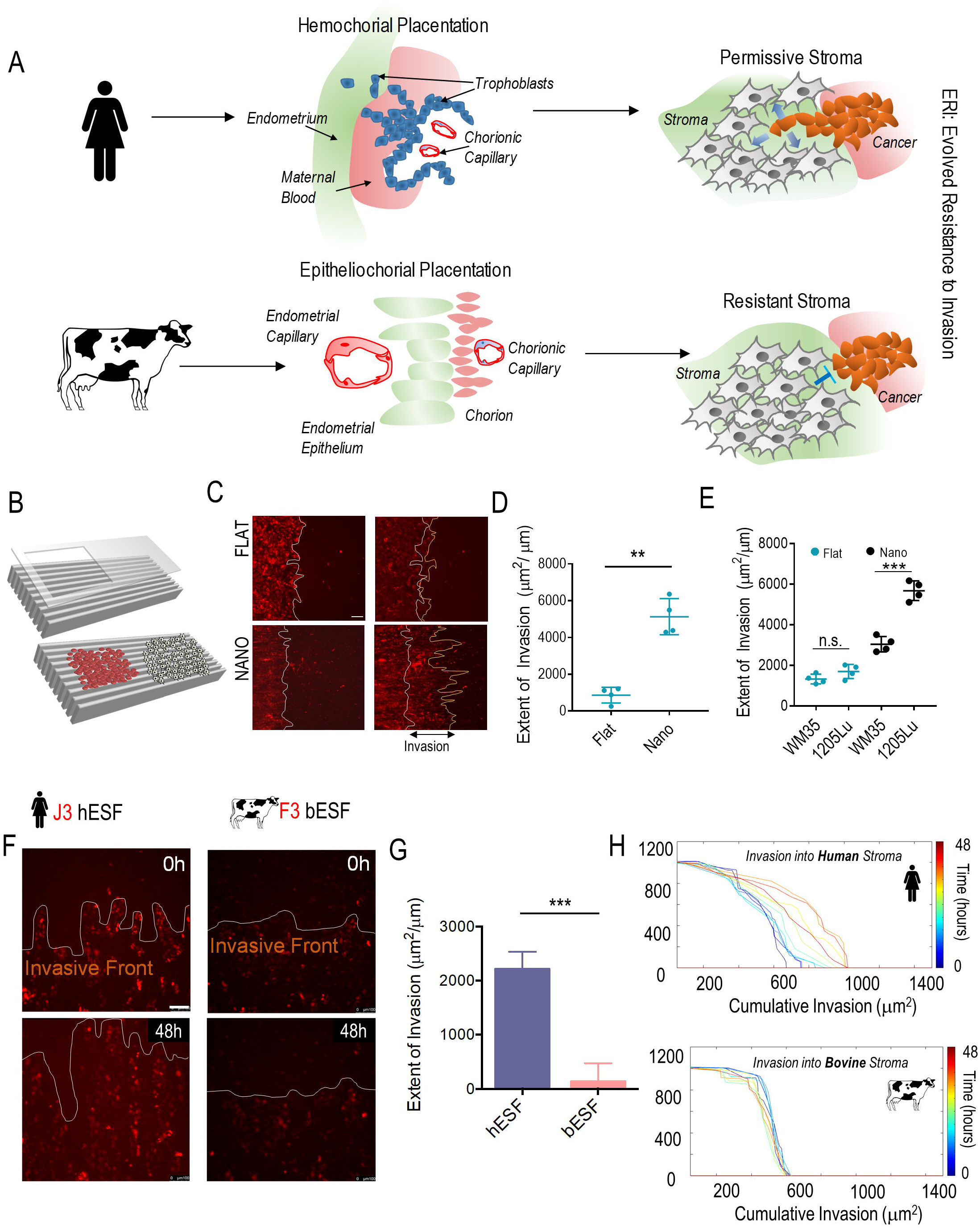
An experimental platform to test the hypothesis of Evolved Levels of Invasibility (ELI). **(A)** Schematic describing the ELI hypothesis. Placentation in humans is hemochorial, wherein the placental trophoblasts invade into the maternal stroma reaching the blood supply. In contrast, in cows and other boroeutherians, placentation has recently evolved to be epitheliochorial, where the trophoblast epithelium attaches to the endometrial epithelium, but does not invade the maternal interstitium. The ELI paradigm states that bovine stroma has evolved to resist invasion compared to human stroma, and therefore secondarily limits cancer metastasis. (B) Schematic showing a cell patterning nanotextured platform to quantitatively and sensitively measure collective invasion into stroma; Stromal cells and invasive cells are patterned by a PDMS stencil into juxtaposed monolayers heterotypically interacting with each other, and imaged using live cell microscopy to observe collective cell invasion into the stroma. (C) Time course images showing invasion of 1205Lu malignant melanoma cells (red) into BJ5ta human skin fibroblasts (unlabeled) for 18 hours on a flat substrate vs a nanotextured substrate; Quantification of the extent of invasion per unit length of heterotypic intercellular interaction shown in (D). (E) Quantification of the extent of invasion of non-malignant WM35 and malignant 1205Lu melanoma into a monolayer of BJ5ta human skin fibroblasts on flat and nanotextured substrata. (F) Time course images showing invasion of human choriocarcinoma derived trophoblasts, J3 (red) into human endometrial stromal fibroblasts (unlabeled); and bovine trophoblasts, F3 (red) into bovine stromal fibroblasts (unlabeled) for 48 hours; See Figure S2 for phase-contrast and Supplementary Movies for dynamics of invasion; Quantification of the extent of invasion of trophoblasts into the respective stromal monolayer shown in (G). (H) Time course dynamic analysis showing cumulative invasion of J3 and F3 into respective species-specific endometrial stromal monolayers. In D, E, and G, n = 4 independent biological replicates; Statistical comparisons made using Student’s t-tests **: p < 0.01, ***: p < 0.001; Error bars denote standard error of the mean (s.e.m.).

## Results

### Collective cell invasion behavior can be modelled in an ECM-mimetic co-culture system

To test the ELI model, we developed a quantitative method permitting measurement of collective cell invasion (CCI) into a lawn of stromal cells *in vitro*. The CCI into stroma is a complex process. Invading epithelial cells migrate while also maintaining cell-cell contacts before a transition to the mesenchymal migration mode would occur^15,16^, create pathways through the stromal mass by mechanical and biochemical separation of stromal cells^17,18^, break cell-cell junctions, and reorganize the ECM^19^. During infiltration and spread, invading cells actively interact with stroma, which likely modulates their invasive capacity^1,20,21^. Importantly, as suggested by *in vivo* imaging of primary tumor dissemination, invasive spread can be triggered and guided by increasingly aligned ECM fibers, serving as ‘tracks’ for contact-guided cell migration away from the primary tumor site^22^. Based on these considerations, we sought to mimic the *in vivo* cell invasion process by reproducing the nanotopographic features of aligned ECM fibers (mimicking their size, anisotropy, and biochemical properties), in combination with a heterotypic cell co-culture. This approach was validated in previous cancer-stroma interaction studies^23,24^. The platform also allowed for sensitive quantitative measurement of collective cell invasion into the stroma compartment (Figure 1C). The extent of invasion was measured as the difference between the final and initial area occupied by the invading cells, and normalized by the initial length of the boundary between the invasive and stromal cells. The resulting measure can be interpreted as the average distance over which the invasive cells penetrated into the stromal cell monolayer.

We first compared the spread of malignant (1205Lu) and non-malignant (WM35) melanoma cells into a monolayer of human skin fibroblasts (BJ5ta) on the ECM-mimicking substratum, or on flat surfaces fabricated of the same material as a control. We found that the nano-textured substratum allowed malignant 1205Lu melanoma cells to more extensively invade into BJ5ta human skin fibroblast monolayer (Figure 1D-E, S1A). Furthermore, when the invasive characteristics of 1205Lu and WM35 cells were compared on a flat surface, no statistically significant difference could be detected within 18 hours of observation and 4 replicates. In contrast, a significant difference between the extent of invasion for 1205Lu and WM35 was clearly detectable on the nano-textured substratum (Figure 1E, S 1B, C), indicating an increased resolution of this assay of collective invasion into stroma.

### Bovine endometrial stroma is resistant to trophoblast invasion

To test the ELI hypothesis, we measured the invasion of a human trophoblast cell line obtained from a choriocarcinoma (J3) into a dense layer of human endometrial stromal fibroblasts (hESFs), and compared it with the invasion of bovine trophoblasts (F3) into a dense layer of bovine endometrial stromal fibroblasts (bESFs). A 48 hour observation showed that J3 cells invaded deeply into hESF cell layer, while the F3 invasion into bESFs was comparatively much more limited (Figure 1F-G, Supplementary Movies). Dynamic analysis of the extent of invasion into the respective stromal models showed that human J3 cells formed invasive forks, which propelled the J3 cell invasion into hESFs, while bovine F3 cell invasion fronts remained stunted with no appreciable formation of invasive forks (Figure 1F, G, H). These results recapitulate the *in vivo* differences observed during embryo implantation, validating our approach to measure trophoblast-stromal interactions^25-28^.

In human placenta, extravillous trophoblasts (EVTs), differentiated from the cytotrophoblasts, constitute the most invasive cell type, displaying a high degree of invasion into the maternal endometrium and even the myometrium^29^ ^30^. We therefore next measured the invasion of human EVT cell line, HTR8, into hESFs, and compared the extent of invasion to that of F3 cells (bovine cytotrophoblasts do not differentiate into EVTs) invading into the layer of bovine stromal fibroblasts (Figure 2A). Our results showed that HTR8 cells invaded deeply into hESFs while F3 did not display appreciable invasion into bESFs over the same time course (Figure 2B). To test whether this difference is due to the identity of the stromal or the trophoblast cells, we also conducted cross-species invasion experiments. We found that bovine F3 trophoblasts displayed a high degree of invasion into hESF, while human trophoblast HTR8 cells invaded bESF to a much lower extent (Figure 2B). These findings indicate that the bovine, but not human ESFs, are resistant to invasion by trophoblast cells of either species. In summary, these results suggest that the degree of trophoblast invasion is, at least in part, controlled by the identity of the stromal cells rather than only by the invasive capacity of trophoblast cells, providing support for the ELI hypothesis.

**Figure 2.**
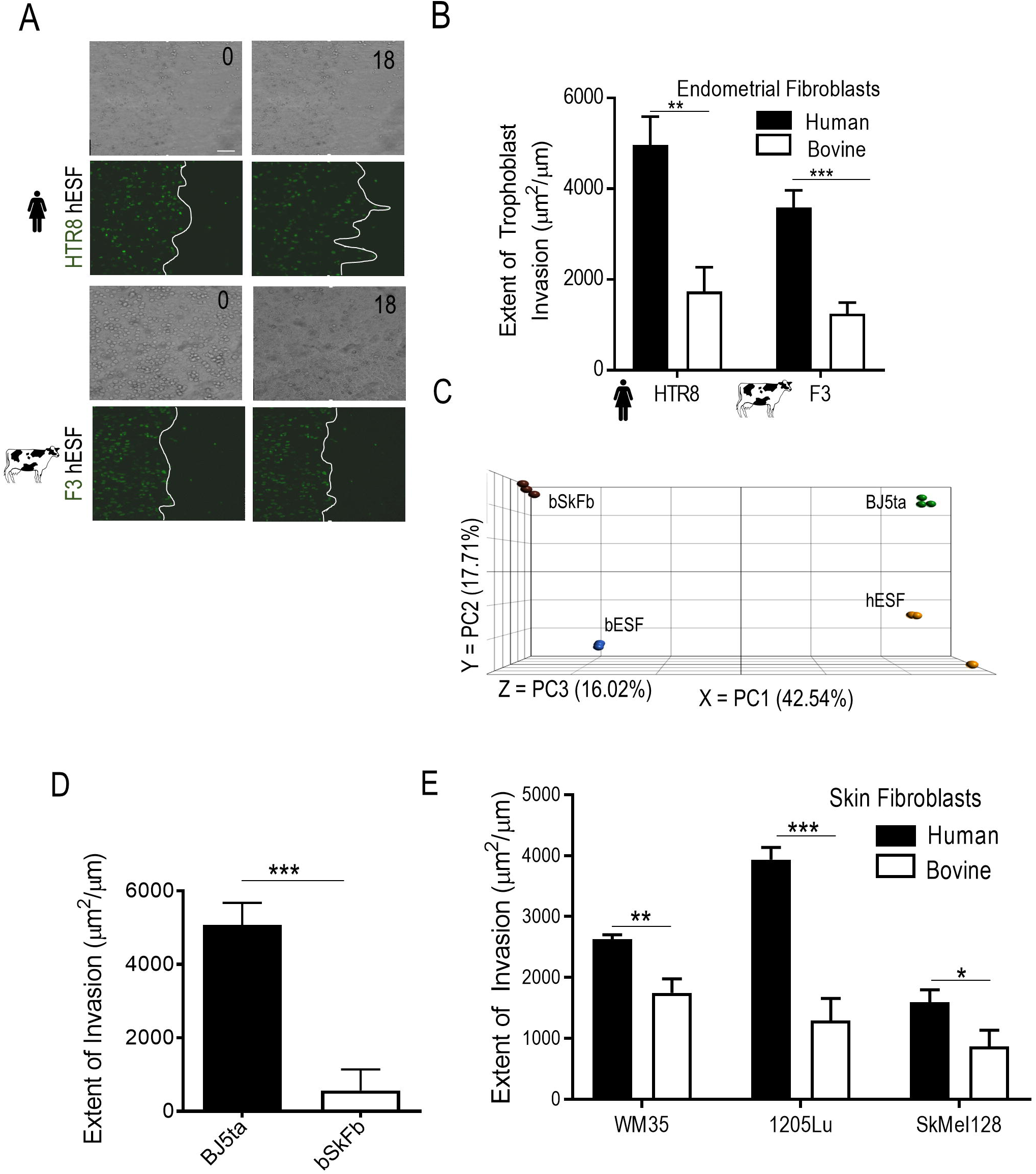
Bovine stroma resists trophoblast and melanoma invasion. (A) Representative time course images of human extravillous trophoblasts HTR8 (green) and bovine trophoblasts F3 (green) invading into endometrial stromal fibroblasts of the respective species for 18 hours; White traces show the boundary of the invasive fronts; Quantification of the extent of invasion for either trophoblasts shown in (B). (C) Principal Component Analysis (PCA) of the gene expression data from skin fibroblasts from human (BJ5ta) and bovine (bSkFb), and endometrial stromal fibroblasts from human (hESF) and bovine (bESF); 3 biological replicates were used for RNA Sequencing. (D) Extent of invasion of malignant A375 cells into BJ5ta and bSkFb monolayers measured over 24 hours. (E) Extent of invasion of other well characterized human melanoma cell lines into BJ5ta and bSkFb monolayers after 18 hours of observation. In B, C, E, n = 3, and in D, n = 8 independent biological replicates; Statistical comparisons made using Student’s t-tests *: p < 0.05, **: p < 0.01, ***: p < 0.001; Error bars denote standard error of the mean (s.e.m.).

### Bovine skin fibroblasts resist melanoma invasion

The findings above support a model postulating that stromal cells from different species resist trophoblast invasion to different degrees, but leave open the possibility that these effects could be specific to the fetal-maternal interface. Would stromal compartments in other tissues, e.g., the skin, show similar species-dependent properties? In humans, the aggressive skin cancer melanoma is an extremely malignant neoplastic disease. Its invasion into the surrounding stroma is a strong prognostic predictor of malignancy^31^ and shares many characteristics with the placental invasion into stroma^32-34^. We thus explored whether gene expression profiles of human and bovine skin fibroblasts are more similar to that of their corresponding endometrial fibroblasts than to each other.

The null hypothesis of independent transcriptome evolution of each cell type predicts that corresponding (homologous) cell types from human and bovine tissues should be more similar to each other than they are to a different cell type in the same species^35^. For instance, one would expect that human and bovine skin fibroblasts are more similar to each other than human skin fibroblasts are to human endometrial fibroblasts. This is because the human and bovine lineages diverged more recently than skin and endometrial fibroblasts have differentiated in evolution. On the other hand, if skin and endometrial fibroblasts co-evolve in each lineage, the opposite pattern is expected, e.g. human skin and endometrial fibroblasts would be more similar to each other than human skin fibroblasts would to bovine skin fibroblasts. The ELI hypothesis predicts that in each species endometrial fibroblasts and other fibroblasts, including in the skin, share the same level of invasion resistance because they co-evolved. Therefore, the ELI hypothesis would predict a pattern where the stromal cell types from the same species are more similar to each other than they are to corresponding stromal cell types in the other species^36^.

To test this prediction, we performed RNA sequencing on isolated bovine skin fibroblasts and immortalized human skin fibroblasts (BJ5ta), and compared the global gene expression changes to human and bovine endometrial fibroblasts. Principal component analysis showed that the genetic distance between skin fibroblasts and ESFs of the same species was less than the distance between corresponding cell types of different species (Figure 2C), suggesting that distinct stromal cell types from the same species have similar expression profiles vs. those of a different species. The observed pattern can be explained by the assumption that endometrial and skin fibroblasts undergo correlated evolution due to pleiotropic effects of mutations on gene expression^35,37^. In other words, these data indicate that the biology of skin and endometrial fibroblasts can be expected to be similar within species.

The transcriptome data suggests that bovine skin fibroblasts might exhibit resistance to invasion similar to that of bovine endometrial fibroblasts. We thus measured the extent of invasion of malignant human melanoma cells (A375) into the skin fibroblasts from human (BJ5ta), and bovine (bSkFb), and found that bSkFb indeed resisted A375 invasion more strongly than human skin fibroblasts (Figure 2D, E). This greater resistance of bSkFb, in comparison to human BJ5ta cells, was supported by a similarly strong resistance to invasion of benign WM35, malignant 1205Lu and SKMel28 melanoma cell lines. A similar assay of invasiveness of bovine melanoma cells was precluded owing to lack of biological material, since melanoma is rarely observed in US-based cow herds as US cows do not reach the age where melanoma would be expected (Dr. Elizabeth W. Uhl, University of Georgia, pers. comm.). Overall, these results indicate that bovine skin fibroblasts, like bovine endometrial stromal fibroblasts, can resist collective cell invasion better than their human counterparts.

### Human and bovine fibroblasts respond differently to trophoblast co-culture

To understand the genetic underpinnings of ELI in bovine stromal cells, we collected RNA sequencing data from endometrial fibroblasts of human and bovine origins, both with and without co-culture with the respective trophoblasts. The particular focus of this analysis was on the genes differentially expressed in either stromal cells, both basally and upon interaction with species-specific trophoblasts. To achieve this, endometrial fibroblasts of both species were labeled with the DiI dye, and co-cultured with equal number of unlabeled trophoblast cells (Figure S3A). Co-cultures were maintained for 72 hours, and the cells were sorted using fluorescence activated cell sorting (FACS), following the method previously used to analyze the genetic correlates of melanoma-endothelial cell co-culture^23^. RNA-Sequencing was performed for RNA isolated from bovine ESFs with and without co-culture with F3 (bESFs and bESF^co-F3^), and for human ESFs with and without co-culture with HTR8 (hESFs and hESF^co-HTR8^).

Analysis of transcript abundance showed a large number of genes differentially expressed in bovine vs. human endometrial fibroblasts (Figure S4B). Human and bovine ESFs responded strongly to co-culture with their cognate trophoblasts cells, as shown in the p-value distributions (Figure S3C-D). Furthermore, human and bovine ESFs responded differently to trophoblast co-culture (Figures 3A-B, Figures S3E-F). For instance, of the 2,135 genes that are upregulated in human ESF in the presence of HTR8 cells (p≤0.01, FDR=3.1×10^−3^) only 349 are also upregulated in the bovine ESF following co-culture. Interestingly, this gene set, shared between human and bovine ESFs, had a 1.87-fold enrichment of genes involved in the regulation of apoptosis (52 genes, p=8.04×10^−6^, FDR=1.53 10^−2^). Human and bovine endometrial stromal cells also differed substantially with respect to the number of genes affected by co-culture, with human cells displaying a larger number of differentially expressed genes (5,349 genes changed at p≤0.01, FDR=3.1×10^−3^) compared to bovine cells (3,101 genes at p≤0.01, FDR=7.01×10^−3^). Gene set analysis further revealed that more genes belonging to chemotactic activity, cell motility, and metastasis ontologies were present at higher relative abundance in hESF^co-H8^ compared to bESF^co-F3^ (Figure 3B). These results suggest that the inter-species difference in resistance of stromal cells to invasion may indeed be genetically encoded in the basal and induced transcriptomes of these cells.

**Figure 3.**
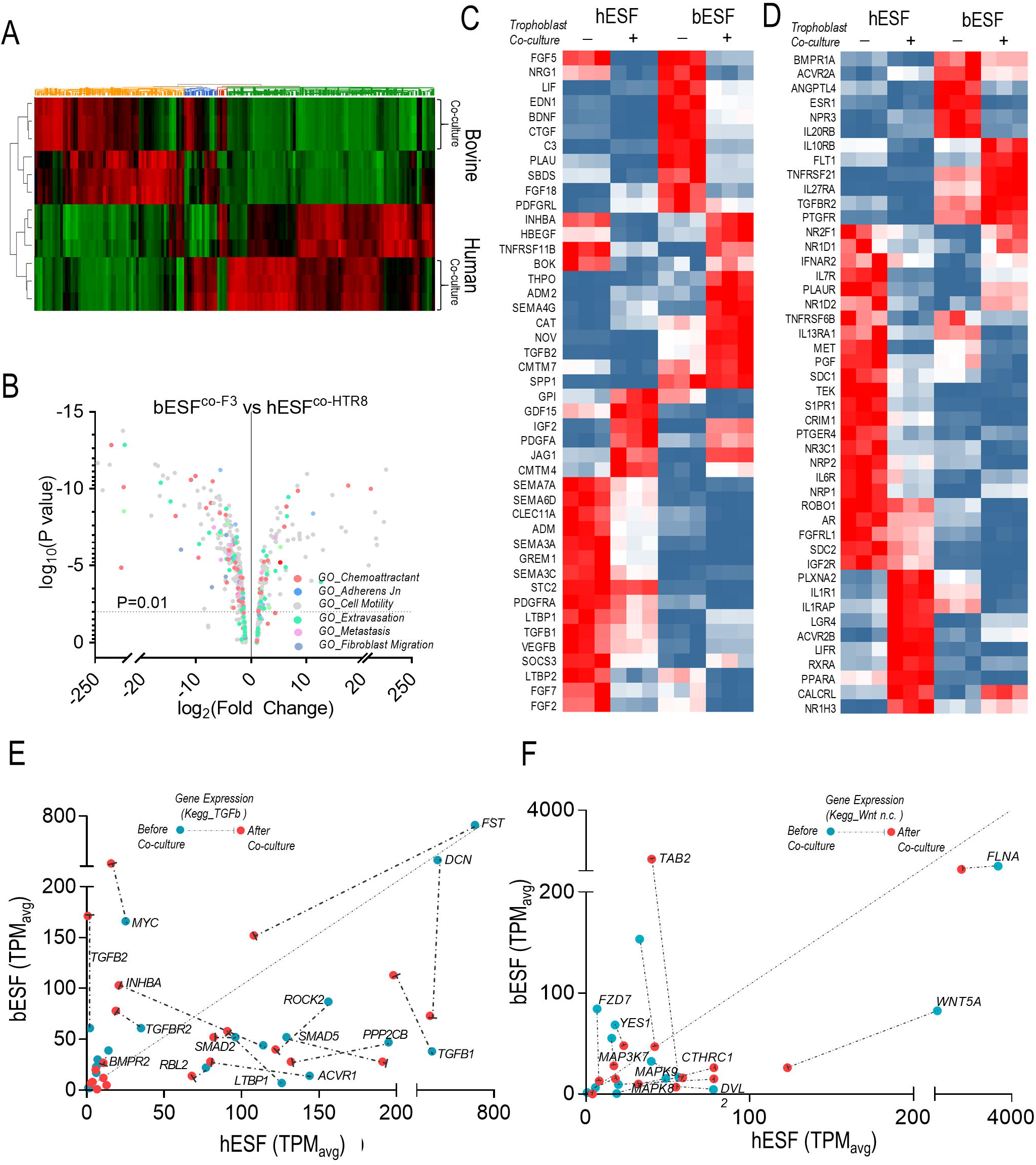
Transcriptomic analysis of bovine and human endometrial stromal fibroblasts reveals differential response to trophoblasts. (A) Heat map of gene expression differences of human and bovine endometrial stromal fibroblasts with and without co-culture with HTR8 and F3 trophoblast cells respectively. (B) Volcano plot showing fold differences in the TPM values of bESFs vs hESFs co-cultured with the respective trophoblasts, color coded for relevant gene ontologies. (C-D) Heatmap showing expression of genes in hESFs and bESFs with or without co-culture with HTR8 and F3 respectively, belonging to the gene-sets chemokine (C) and chemokine receptors (D). TPM values of genes significantly different in hESFs and bESFs for KEGG pathways for TGFB (E), and non-cannonical WNT signaling (F); For each gene, TPM values before and after co-culture are shown in green, and red dots respectively; Arrows show the change in TPM values due to trophoblast co-culture; Also are shown coefficient of determination, R^2^ for the linear regression of TPM values between hESF and bESF, and the standard deviation of the residuals, Sy.x.

We then more fully explored the effect of co-culture on gene ontology (GO) enrichment to examine putative mechanisms of bESFs resistance to trophoblast invasion. We found that human and bovine ESFs expressed markedly different sets of chemokine ligands (Figure 3C) and chemokine receptors (Figure 3D), although, interestingly, both were enriched in transcripts associated with endothelial cells and angiogenesis. Genes showing elevated expression in hESFs included Fibroblast Growth Factors (FGFs), Vascular Endothelial Growth Factors (VEGFs), Semaphorins, members of the Transforming Growth Factor (TGF) family, as well as NRPs and *ROBO1*, whereas bESFs showed high expression of Endothelin (*END1*), Plasminogen activator (*PLAU*), Thrombopoeitin (*THPO*) and notably, Transforming Growth Factor beta 2 (*TGFB2*). In contrast, genes belonging to ontologies GO_Adherens Junction and Endothelial Barrier did not show systematic expression differences in favor of either hESF or bESF (Figure S4A-B). Finally, the genes related to GO_Fibroblast Migration tend to be expressed at lower levels in bESF vs. hESF (Figure S5C). These data point to the possibility that stromal response to trophoblast co-culture varies strongly between the species for genes that could play a critical role in the interaction between invading and stromal cells, opposed to genes which regulate basal stromal integrity. Differential expression of these genes may affect the invasibility phenotype in either species.

These basic analyses hinted at a possibility that molecular interactions between the stromal and invading cells were drivers of ELI in human and bovine. Molecules in the Transforming Growth Factor (TGFB), and WNT signaling pathways have been key mediators of cancer stromal interaction in various tissue types^38-43^. Ingenuity pathway analysis also showed that both signaling pathways were differentially activated in bovine and human ESFs, consistent with the results above (Figure S5E). We therefore explored the effect of co-culture of respective trophoblasts on the activation of these signaling pathways in either species. Expression of genes in non-canonical WNT and TGFB pathways were indeed higher in hESFs compared with bESFs (Figure 3E, F). Although co-culture with trophoblasts resulted in reduction in expression of genes belonging to TGFB pathway for hESFs, they continued to remain many-folds higher compared in bESFs. While TGF ligands TGFB1, and TGFB3, as well as receptors of the TGFB family, e.g. ACVR1 were highly expressed in hESFs, negative regulators like TGFβ2 and inhibin A were expressed at higher levels in bESFs. Similarly, we observed that downstream transcription factors Smad 2, −3 and −5, as well as the co-SMAD, SMAD4 were expressed at higher levels in hESFs compared to bESFs (Figure 3E). Similarly, many members of the WNT signaling network displayed higher expression in hESF vs bESF (Figure 3F, S5D). These results supported a role for paracrine signaling between heterotypic cells during the invasion processes, prompting us to further explore and validate the role of the signaling networks in the stromal invasion.

### Selected gene knockdown in human fibroblasts increases resistance to melanoma invasion

We showed that bovine endometrial fibroblasts, as well as bovine skin fibroblasts, are more resistant to invasion compared to their human counterparts. Transcriptomic analysis revealed large differences in gene expression, and differential responses to interaction with trophoblasts of either species. We therefore hypothesized that modulating the differentially expressed genes in human stromal cells could induce them to become more similar to their bovine counterparts in displaying an increased resistance to invasion in human cells. Given our results, we focused on genes related to WNT and TGFB signaling, and those interacting with these pathways. We further limited the gene set to those genes whose expression was affected in response to trophoblasts co-culture in either species. Specifically, we selected 16 genes from the TGFB and WNT pathways that had higher expression in hESF or were upregulated in hESF in co-culture with trophoblast cells (Figure 4A). We then targeted these genes using a battery of siRNAs in hESFs and BJ5ta to test whether they could modulate stromal invasibility.

**Figure 4.**
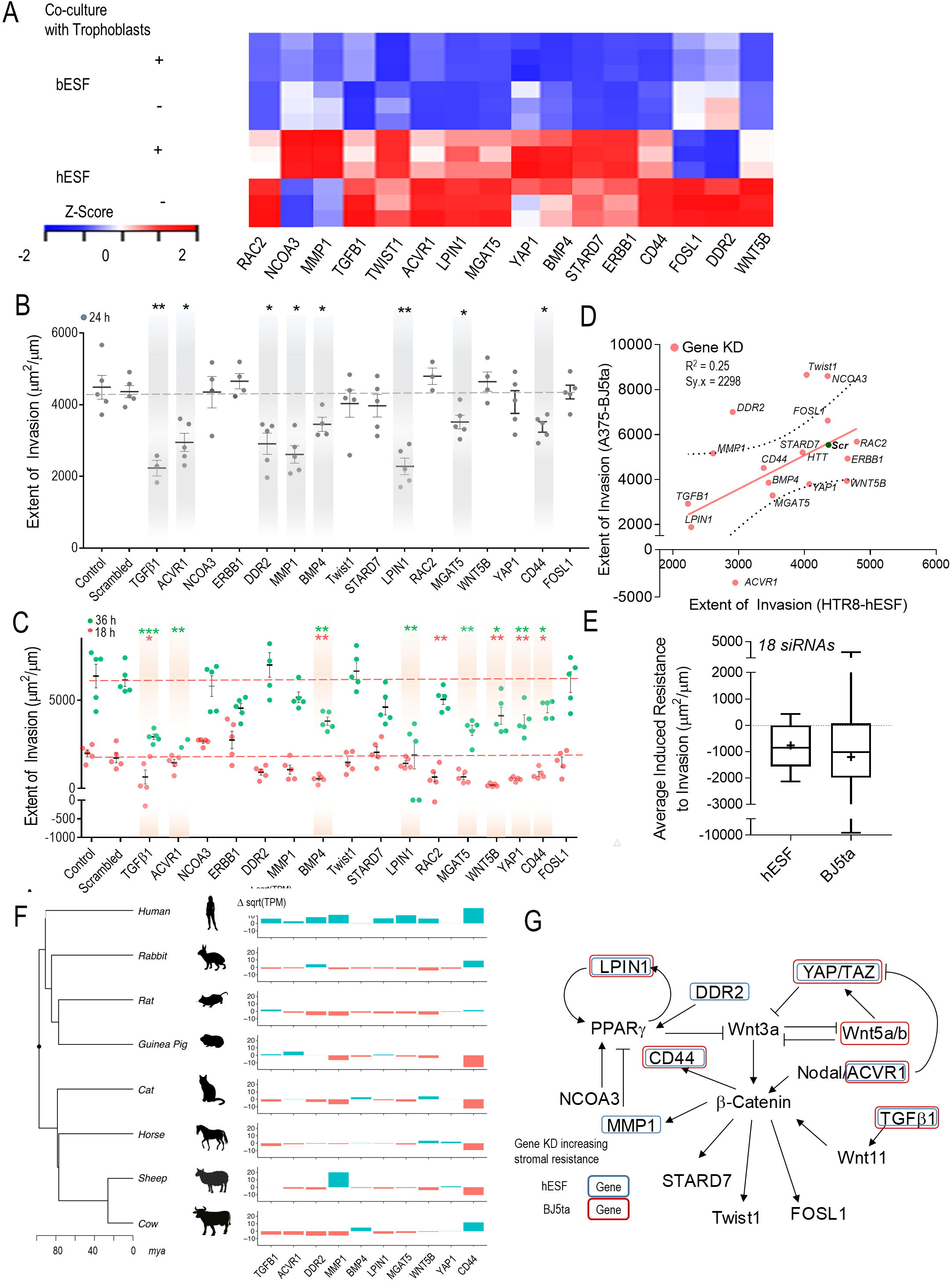
Induced resistance to invasion in human stroma by evolutionarily inspired gene silencing. (A) Heatmap showing selected set of genes used for gene silencing in human stromal fibroblasts; Shown are z-score of TPM values for each transcript. (B-E) siRNA based gene knockdown in human stroma to induce expression similar to bESF increases resistance to invasion; (B) Extent of invasion of HTR8 cells into hESFs subjected to siRNA mediated gene silencing measured over 24 hours using nanotextured platform; (C) Extent of invasion measured for 18 hours (red), and 36 hours (green) using nanotextured platform for A375 invasion into BJ5ta subjected to siRNA induced silencing of individual genes; Each dot in B, C reflects an individual invasion observation, Error bars show s.e.m., Statistical comparisons made with scrambled siRNA using Student-t-test; *: P < 0.05, **: P < 0.01, ***: P < .0001. (D) Correlation shown between the extent of invasion of HTR8 into hESF, and A375 into BJ5ta for each siRNA based gene silencing; (E) Average extent of invasion in hESF and BJ5ta for the above 20 gene silencing normalized to the extent of invasion observed for control scrambled siRNA knockdown in either human stromal cells; Error bars show min to max values; Average values corresponding to each siRNA knockdown shown as individual dots. (F) Relative transcript levels of the selected genes in various boroeutherians species compared to their most common ancestor; red colored bars denote decrease, and green bars show difference in sqrt(TPM) values vs the common ancestor; mya refers to million years ago. (G) Network of WNT signaling and its interaction with TGFB pathway showing previously known interactions, mapped with the gene silencing induced resistance in hESF, and BJ5ta stromal cells.

Comparison of each siRNA-transfected stromal cell population was made with appropriate untransfected controls, as well as with cells transfected with control non-targeting siRNA (Figure S5A-B). HTR8 invasion into hESF was significantly reduced in eight out of 16 genes after 24 hours of observation (Figure 4B), with other knockdowns showing a more limited effect. A375 invasion into BJ5ta skin fibroblast monolayer was observed for 18 hours and 36 hours after gene knockdowns were performed. In this case we found nine gene knockdowns significantly decreasing invasion (Figure 4C). These included members of WNT superfamily, TGFB ligands, as wells as less established targets and effectors of WNT signaling, e.g. STARD7^44^, LPIN1^45-49^, and YAP1^50^. Furthermore, we detected a weak but significant correlation in the increase of stromal resistance to invasion following gene knockdowns in both hESF and BJ5ta (r^2^=0.25, p = 0.02), indicating that gene silencing enhances resistance to invasion in a similar manner in both human endometrial and skin fibroblasts (Figure 4D). Moreover, the average response for all experimental gene knockdowns was an increase in resistance to both HTR8 and A375 invasion (Figure 4E). These results experimentally validated the putative role of these genes in regulating stromal invasibility, and also suggested that human skin fibroblasts could indeed be induced to resist melanoma invasion.

### Opposite evolutionary trends of vulnerability to malignancy in humans and bovines

In order to assess the evolutionary history of the genes identified above as causal in enhancing invasibility of fibroblasts, we isolated and cultured the endometrial stromal fibroblasts from rabbit, rat, guinea pig, cat, horse and sheep to extend our analysis beyond those of human and cow (Figure 4F). This phylogeny represents two clades of eutherian mammals, the Euarchontoglires, largely composed of rodents and primates, and the Laurasiatheria, largely composed of ungulates and carnivores. These two clades together form the clade of Boreoeutheria. We plotted the phylogenetic tree of these eight species based on the tree by Meredith and coworker^51^, along with the expression of these genes in skin fibroblasts compared to the inferred boreoeutherian ancestor 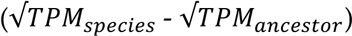. We then reconstructed the gene expression changes over the phylogeny for 10 genes that have been found to influence invasibility in our knockdown experiments.

We indeed found that bovines evolved lower expression of many genes that enhance invasibility (*TGFB1, ACVR1, DDR2, MGAT5, MMP1*), confirming the hypothesis that the identified genes which enhance invasiveness have evolved lower levels of expression in the bovine lineage coincidental with the evolution of secondarily non-invasive placenta in bovines and related animals. Surprisingly, humans evolved higher expression levels of invasibility enhancing genes (*TGFB1, ACVR1, DDR2, LPIN1, CD44, MMP1*), compared to the inferred boreotherian ancestor. Thus, the inference initially suggested by the apparent divergence in evolutionary trajectories of the primate and the bovine lineages based on differentially expressed gene sets, was further supported by the finding that the expression levels of these genes in human skin fibroblasts is higher than that of rabbit, rat and guinea pig which are all included in the clade of Euarchontoglires. All of these species have hemochorial placentation and thus the higher expression level of these genes in humans likely evolved in the primate lineage. Further, our taxon sample is balanced with four species from Euarchontoglira (human, rabbit, rat and guinea pig, all with hemochorial placenta) and Laurasiatheria (cat, horse, sheep and cow, all with less invasive placentae, either endotheliochorial or epitheliochorial placenta). The limited data on a selected group of genes, therefore, supports the hypothesis of Evolved Levels of Invasibility (ELI) amongst mammals, but points to the possibility of increased stromal vulnerability in humans, and increased stromal resistance in the ungulates.

## Discussion

Mammalian species differ in their potential for tumorigenesis, as well as their vulnerability to cancer metastasis^7,52^. The comparative biology of cancer across different animals has identified unique tumor suppressor mechanisms explaining the variation in occurrence of tumors between species. For instance, the naked mole rat is extremely long lived among rodents, and has evolved at least five different tumor suppressor mechanisms^53,54^. Large mammals such as elephants have lower cancer occurrence than expected from their size and life span, an observation referred to as the Peto’s Paradox^55,56^, in part, because of evolutionary changes to the TP53 gene family^57,58^.

Here, we focus on differences across species of the rates of cancer malignancy rather than differences in rates of tumorigenesis. For instance, melanoma occurrence is common in older bovines and equines, but remains largely benign^59-61^; while it is highly malignant in humans. This correlates with the phenotype of the fetal-maternal interface, i.e. the degree of placental invasion during pregnancy. The results presented here show that this species difference may, in part, be caused by species differences in invasibility of stromal cells, both in the uterus as well as the skin. Comparative transcriptomics showed large differences in gene expression between cow and human stromal cells. Our data suggests that bovine stroma evolved lower invasibility via decreased paracrine signaling through TGFB and WNT pathways. In particular, these results argue that TGFB secretion, and high non-canonical WNT signaling in stromal cells are causal factors explaining the high vulnerability of human stromal tissues to cancer invasion, at least in the case of melanoma. Comparative transcriptomic data across multiple additional species further suggests that the human lineage evolved higher expression of these genes enhancing tumor and trophoblast invasion and thus likely have evolved higher malignancy rates than were present in the common ancestor of eutherian mammals.

siRNA-guided knockdown in human stromal fibroblasts revealed genes that impart stromal invasibility by melanoma and trophoblast cells (Figure 4G). These genes include those encoding secreted TGFB ligands, consistent with results from colorectal cancer showing that stromal expression of these ligands predicts poor prognosis^41^. Other genes enhancing stromal invasibility are members of the non-canonical WNT pathway, as well as genes that are known to modulate WNT signaling via PPARG activity, e.g. *LPIN1*^62^, by directly regulating β-Catenin levels, e.g. *YAP/TAZ*^50^, or by being regulated by β-catenin induced transcription, e.g. *CD44*^63^. Of these, YAP1, a Hippo pathway target promoting tumor growth^64^, can also be regulated by non-canonical WNT signaling, and can in turn inhibit canonical WNT signaling^50^. PPARG a regulator of lipid metabolism is involved in various cancers, although the evidence is mixed about its role in driving cancer metastasis^65^. *LPIN1*, a negative regulator of PPARG activity, can therefore also be linked to Wnt signaling through this regulatory molecule. Both TGFB^66,67^ and non-canonical WNT signaling^68,69^ are known to affect tumor progression and metastasis. TGFB1 and TGFB3, expressed at relatively higher levels in hESF, are also reported to regulate β-catenin activity, indicating that human stromal vulnerability to invasion may be influenced by paracrine signaling to invading cells through secreted ligands. We identified differential expression of not only WNT and TGFB family members, but also other ligand families that might potentially be involved in interaction between invading and invaded cell populations. Also of interest, many of the genes identified in our comparative gene expression screen have connections to metabolic regulation. These include genes encoding LIPIN1 which regulates triglyceride metabolism, ^70^ and MGAT5, which encodes N-acetylglucosaminyltransferase V, known to regulate glucose uptake in tumor cells^71^. Our analysis indicates that these genes may be crucial in regulating stroma-assisted invasion, and that the stroma of hoofed animals has evolved lower expression levels of these genes with the effect of decreasing invasibility by trophoblasts.

Our data support the ELI hypothesis, suggesting that differences in stromal gene expression between species are causal in determining both embryo implantation as well as stromal resistance to early cancer dissemination. Epitheliochorial placentation has evolved multiple times from the pre-existing invasive placentation, suggesting evolutionary advantages to non-invasive placentation. Epitheliochorial placentation may be more advantageous for long pregnancies in large animals, where maternal control over the allocation of resources during pregnancy is higher^72^. In contrast, the more invasive hemochorial placentation is believed to be a fetal advantage, with increased transfer of nutrients across the maternal-fetal interface, and is associated with higher fetal growth rate^73^. Indeed, hemochorial placentation may be correlated with increased fetal brain growth^74,75^, and according to our hypothesis, may be an evolutionary compromise resulting in correlatively higher vulnerability to cancer dissemination. Although we have looked at a small subset of genes differently expressed between bovine and human stroma, the expression pattern of these selected genes suggest an opposite trend in humans compared to bovines. Our hypothesis predicts that as many species (including the bovines) increased their stromal resistance to invasion, apes may also have evolved to increase stromal receptivity to invasion, and correlatively have evolved higher vulnerability to cancer malignancy. As extravillous trophoblasts have evolved recently in great apes, the stroma may also have increased invasibility to accommodate the more invasive trophoblast types^76^. Our hypothesis, and its experimental validation, suggests the genetic basis for evolved resistance to invasion can identify relevant causal factors determining the stromal control of cancer progression and provide genetic information to rationally identify novel therapeutic targets.

## Supplementary Figures

**Supplementary Figure 1. Nanotextured stromal invasion assay quantitatively and sensitively measures collective intrusion of cells into stroma. (**A) Time stamped images at 0 and 18 hours showing extent of invasion by DiI labeled 1205Lu cells (red) into BJ5ta stromal fibroblast monolayer (unlabeled) on flat and nanotextured substrata; Quantification shown in Fig. 1D. (B-C) Sensitive measurement of differences in invasion by nanotextured stromal invasion platform; Time stamped images at 0, 6, 12, and 18 hours showing extent of invasion by DiI labeled non-malignant WM35 (B), and malignant 1205Lu (C) cells into BJ5ta monolayer shown in both flat, and nanotextured cell patterned assay; Quantification shown in Fig. 1E.

**Supplementary Figure 2. Human and bovine trophoblasts exhibit different invasive potential into endometrial stroma.** Time stamped images showing extent of invasion of human choriocarcinoma derived trophoblasts, J3 (red) (A), and bovine trophoblasts, F3 (red) (B) into their respective endometrial stromal fibroblast monolayers at 0, and 48 hours.

**Supplementary Figure 3. Human and bovine endometrial stromal fibroblasts respond differently to co-culture with trophoblasts. (**A) Experimental plan to isolate endometrial cells after co-culture with respective trophoblast; (B) Volcano plot showing fold change in genes between bovine vs human endometrial stromal fibroblasts, along with their significance depicted in p value. (C) P-value distribution of t-tests comparing human ESFs with and without co-culture with HTR8 trophoblast cells. P-value distribution of t-tests comparing bovine ESFs with and without co-culture with F3 trophoblast cells. In both C and D, note that thin right hand tail of the distribution, which indicates that a large number of genes are differentially expressed in response to the presence of the corresponding trophoblast cells. (E-F) Scatter plots showing relative TPM values of genes between (E) hESF and hESF co-cultured with HTR8, and (F) bESF and bESF co-cultured with F3; Red dots refer to individual gene transcripts abundance significantly different in between the compared conditions.

**Supplementary Figure 4. Human and bovine endometrial stromal fibroblasts do not appear to differentially respond to their respective trophoblasts by regulating cell-cell adhesion.** Gene ontology analysis of individual genes belonging to GO_Adherens (A), GO_Endothelial Barrier (B), GO_Fibroblast Migration (C), and Kegg ontology for Wnt Signaling (D) for hESF and bESF shown with their relative TPM values; Also are shown the coefficient of determinant, R^2^ for the linear regression between TPM values of hESF and bESF in a given gene-sets, along with the standard deviation of the residuals, Sy.x. Ingenuity Pathway Analysis of selected signaling pathways differentially activated in hESF and bESF; Shown are –log(p-values) of genes in the respective pathways differentially activated, while the color depicts the z-score.

**Supplementary Figure 5. siRNA induced gene knockdown reduces transcript levels in hESFs and BJ5ta cells.** qRTPCR results showing the percentage knockdown of transcript levels in hESF (A) and BJ5ta (B), calculated using ΔΔCt method, compared using GAPDH as housekeeping gene with scrambled siRNA as control.

### Materials and Methods

#### Cell culture

Human ESFs were grown in phenol-red free DMEM/F12 with high glucose (25 mM), supplemented with 10% charcoal stripped calf serum (Hyclone), and 1% antibiotic/antimycotic (Gibco). Decidualization was induced in ESFs with 0.5 mM 8-Br-cAMP (Sigma), and 0.5 mM of progesterone analog medroxy-progesterone acetate (MPA) for 96 hours in DMEM supplemented with 2% charcoal-stripped calf serum (Hyclone). BJ5ta (ATCC) cells were cultured in 80% DMEM and 20% MEM supplemented with 10% FBS, 1% antibiotic/antimycotic, and 0.01 mg/ml hygromycin. F3 cells were obtained from Dr. Pfarrer’s group, and were cultured in DMEM with high glucose, supplemented with 10% FBS, 1% antibiotic/antimycotic (Gibco). J3 cells were cultured in αMEM supplemented with 10% FBS, 1% antibiotic/antimycotic (Gibco), while HTR8/SVNeo (ATCC) were cultured in RPMI-1640 supplemented with 10% FBS, and 1% antibiotic/antimycotic (Gibco).

#### Fabrication of poly-urethane acrylate mold

Photoresist was spin coated on silicon wafers, and electron-beam lithography was used to nanopattern the wafers (JBX-9300FS, JEOL). After the photoresist was developed, exposed silicon was etched by deep-reactive ion etcher (STS ICP Etcher) resulting in formation of sub-micron parallel ridges. Residual photoresist was removed using ashing, and diced into silica masters for subsequent replica molding. UV curable poly-urethane (PUA) was drop-wise dispensed onto the silicon master previously prepared, and contacted with polyethylene terephthalate (PET) film. Application of UV (*v*= 200-400 nm, 100 mJ/cm^2^) for 1 minute was used to cure PUA, after which the mold was peeled off with a pair of tweezers and cured overnight under UV to terminate the residual acrylate groups. The process resulted in PUA mold of ∼50 μm thickness.

#### Fabrication of nanotextured substrate

Previously prepared PET mold was used as a replica mold to transfer the topographic pattern on glass substrate using the technique of capillary force lithography. Briefly, glass substrate was cleaned using NaOH (0.1 M for 1 hour), washed with DIH_2_O, and dried under nitrogen stream. Primer (phosphoric acrylate and propylene glycol monomethyl ether acetate in a ratio of 1:10) was spin-coated on the coverslip as a thin layer, and baked for 20-30 minutes at 70°C. 150 μl of PEG-DA (molecular weight = 575) precursor was dispensed dropwise on the primed cover slip and the PET mold was placed reversibly. After the PUA precursor filled the submicron cavities by capillary action, the substrate was cured in UV (*v*= 250-400 nm, 100 mJ/cm^2^) for 1 minute. The mold was peeled after polymerization using a pair of tweezers. The substrate resultant from the method consisted of nanotextured grooves with an expected elastic modulus of ∼ 70 MPa. The substrate was cured again for an hour under UV to terminate residual active acrylate groups.

#### Cell patterning for stromal invasion assay

For cell patterning, we used a stereolithographic plastic mold to create a polydimethylsiloxane (PDMS) stencil. PDMS stencil was cast by mixing monomer and cross-linker in a ratio of 10:1, and cured at 80oC for 4 hours by placing on a pre-designed plastic mold. Stencils were washed using isopropyl alcohol and dried using N_2_ stream, placed on the nanotopographic substrate coated with laminin (25 μg/ml), and the device kept in vacuum to allow air under the stencil to be removed to avoid chance of leakage. DiI-labeled invasive cells were seeded on the stencil at a density of 10^6^ cells/100μl, and allowed to attach and polarize for 8 hours. Unattached cells were washed off twice with PBS, and the stencil removed carefully using a pair of blunt-end tweezers. Unlabeled stromal cells were seeded at a density of 10^6^ cells/100μl to attach to the area previously covered by PDMS stencil. After 6 hours of attachment, unattached cells were washed off, and the substrate mounted for live cell microscopy. To ensure selection of definitive fibroblasts, and to avoid selection of trophoblasts taking up dye from dead stromal cells and occasional heterotypic cell fusion events, we only collected DiI^high^ cells. All experiments were conducted in similar culture conditions, by 1:1 mixture of the media in either comparable conditions.

#### Live Cell Fluorescence Microscopy

Phase-contrast and epifluorescence microscopy was performed using Leica DMi8 model, with an HC PL Fluotar 10x/0.30 Dry objective, and images were acquired using Leica LAS X Software, and processed using Fiji Image Analysis Software. Lumencor SpectraX was used as a source of light for excitation, and routed through a Rhodamine excitation filter cube consisting of Excitation 546 nm/ 10 nm (band-pass filter), Dichroic 560 nm (long-pass filter), and Emission 585 nm/ 40 nm (band-pass filter), and acquired using Andor EMCCD iXon Ultra 888 camera.

#### Invasion Analysis

Acquired sequential images for each condition were analyzed using Fiji Image Anlaysis Software after contrast enhancement. DiI labeled invasive cell fronts (trophoblasts or melanoma cells) were identified manually for each time point. Area occupied by DiI^+ve^ cells was measured at each time point. Total invasion was calculated as follows:

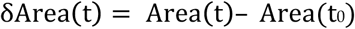

Extent of invasion per unit measurement of interface was determined by dividing δArea by the length of initial intercellular interface

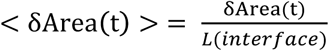

#### Fluorescence Assisted Flow Sorting

Cells were detached from the substrate using TrypLE solution (Gibco), quenched with excess medium, and washed thrice with phosphate-buffered saline (PBS). Isolated cells were suspended in 1% AlbuMAX (Gibco) dissolved in PBS and sorted using fluorescence assisted flow sorting (FACS). FACS was performed using BD FacsARIA II using PE.Cy5 channel, and analysis performed using FACSDiva 6.0. In order to increase the purity of ESFs collected after the co-culture experiments, and to account for possible uptake of dye from dead or dying stromal cells by the trophoblasts, as well as for occasional cell-fusion events, we only sorted DiI^hi^ cells. Sorted cells were collected directly in RNASelect to limit RNA degradation. Even cells that were not in co-culture were subjected to the same sorting protocol.

#### siRNA Transfection and Characterization

For gene silencing experiments, stromal cells (BJ5ta, or hESFs) were cultured in 12 well plates transfected with at least two siRNAs (shown in table below) at a concentration of 50 nmoles per well. Lipofactamine RNAiMAX was used to transfect siRNAs, obtained from IDT.

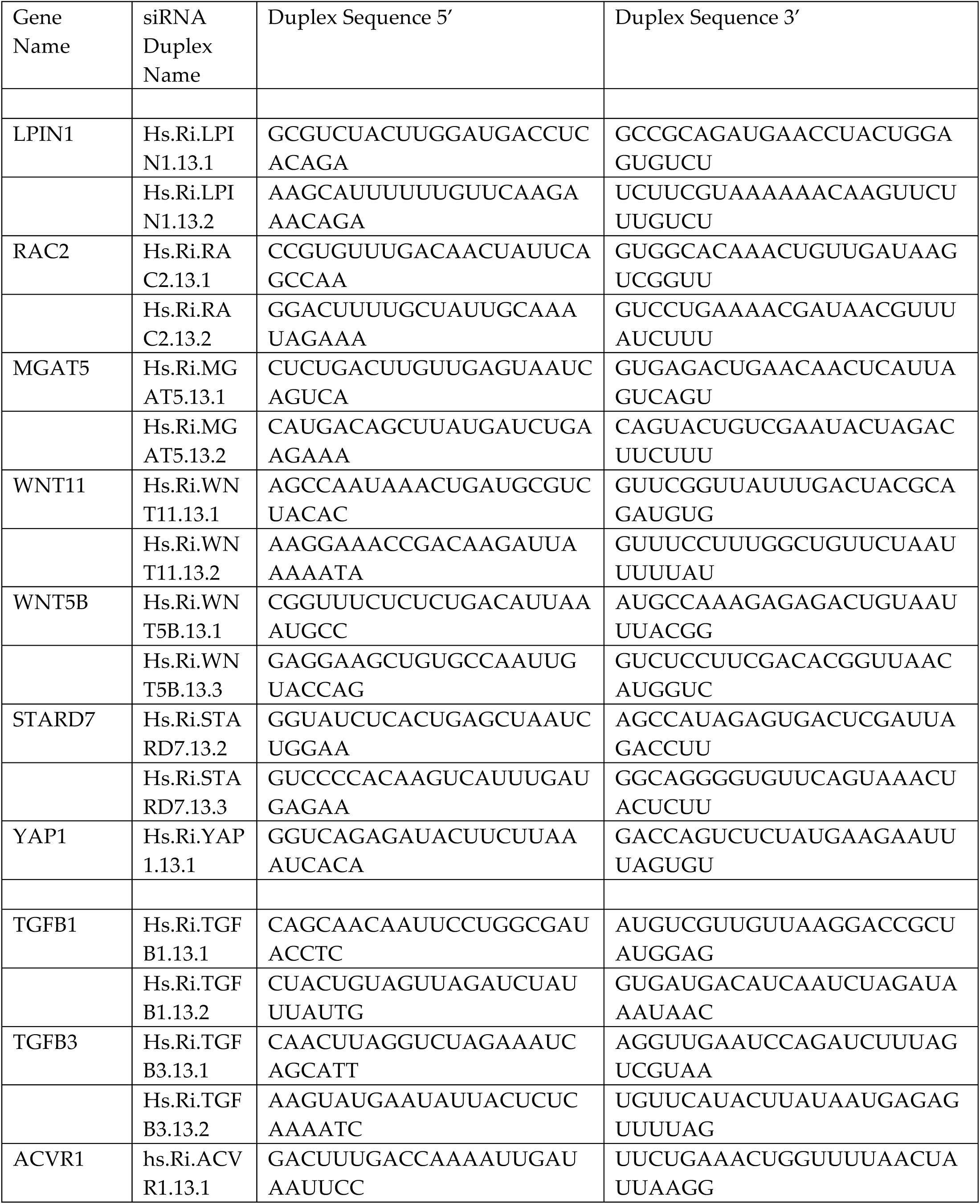

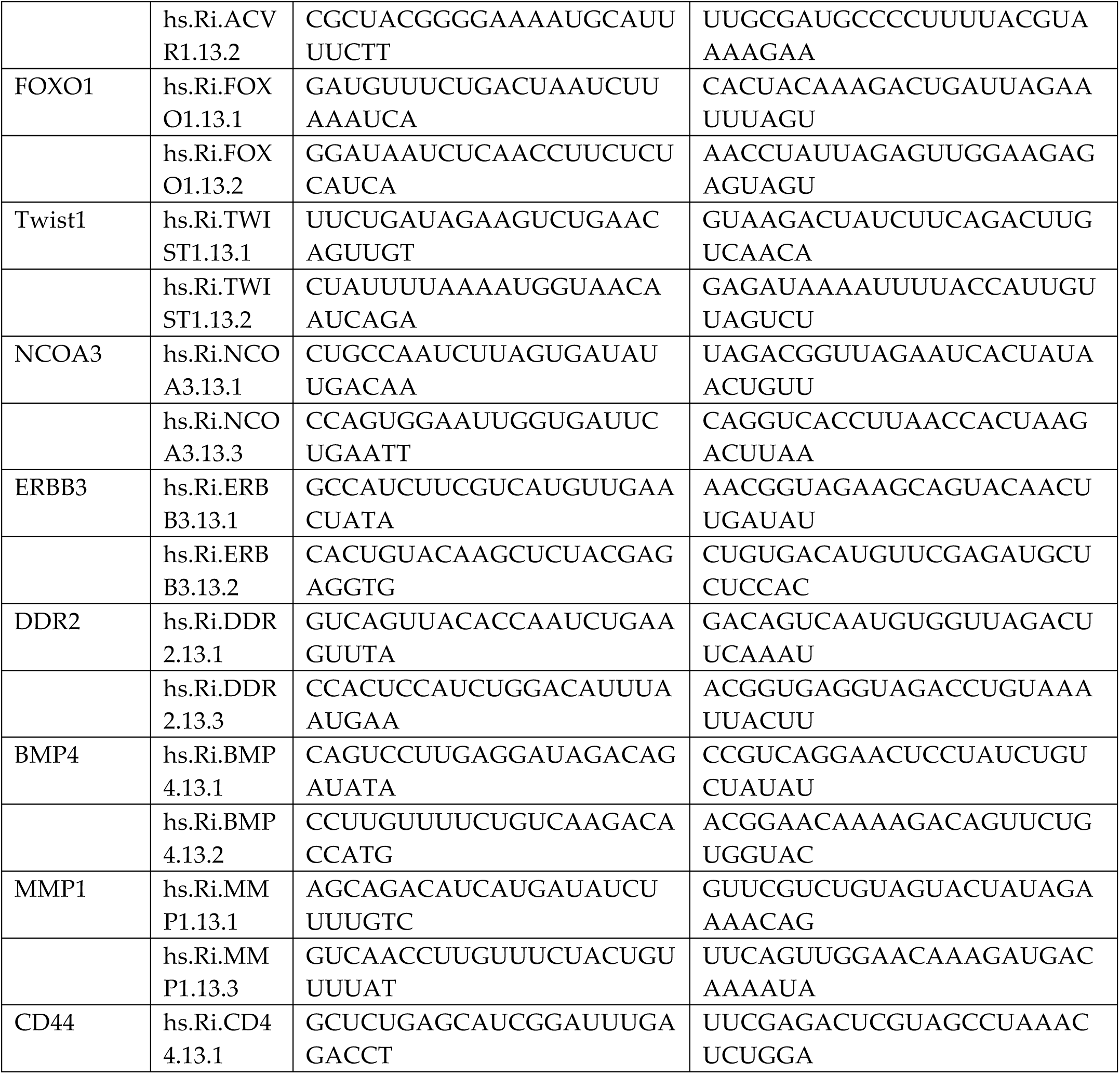

#### RNA isolation and qRT-PCR

RNA was isolated using RNeasy micro kit (QIAGEN, Germany) and resuspended in 10 µL water. SuperScript IV VILO Master Mix (ThermoFisher, USA) was used to synthesize cDNA from 1µg mRNA and ezDNase was used to remove DNA after reverse transcription. Primers from Integrated DNA technologies (IDT, USA) were used for qRT-PCR reaction using SybrGreen (ThermoFisher, USA) to evaluate the efficiency of siRNA mediated knockdown. B2M and RPL0 were used as internal reference genes, and both were found to be consistent within samples after knockdown. qRT-PCR was performed in Quantstudio3 (ThermoFisher, USA) and relative quantitation was performed by comparative C_t_.

#### RNA Sequencing

Agilent Bioanalyzer 2100 was used to determine RNA quality and RIN number of over 8 was observed in the samples. TruSeq RNA Library from illumina was used to prepare mRNA library. These libraries were sequenced on illumina HiSeq to generate 30-40million reads per sample (Single end 75 bp reads) and a high Q score was observed (Q>30) for the sequenced data. Alignment of reads and gene specific analysis was performed in Partek® Flow® software, version 5.0 Copyright ©; 2018 Partek Inc., St. Louis, MO, USA. Human cells data was aligned to reference genome (hg38) using STAR alignment (in Partek Flow) and bovine data was aligned using Bos Taurus assembly UMD_3.1.1/bosTau8. Quantification was performed using quantify to annotation model (Partek E/M) using Ensemble Transcripts release 85 (humans) or Ensemble-UMD3.1 gene annotation model. Normalization was performed using TPM (Transcripts Per Kilobase Million^77^). Gene level comparison and statistical analysis was performed using ANOVA and significance of (adjusted) p0.05 or less was considered for analysis. Gene lists comparison was performed by obtaining Ontology/pathway specific list from Gene Ontology (GO^78^) or kyoto encyclopedia of genes and genomes(KEGG^79^)

