## Supplementary figures and images for "The Evolution of Placental Invasion and Cancer Metastasis are Causally Linked"

### Suppl Figure 1

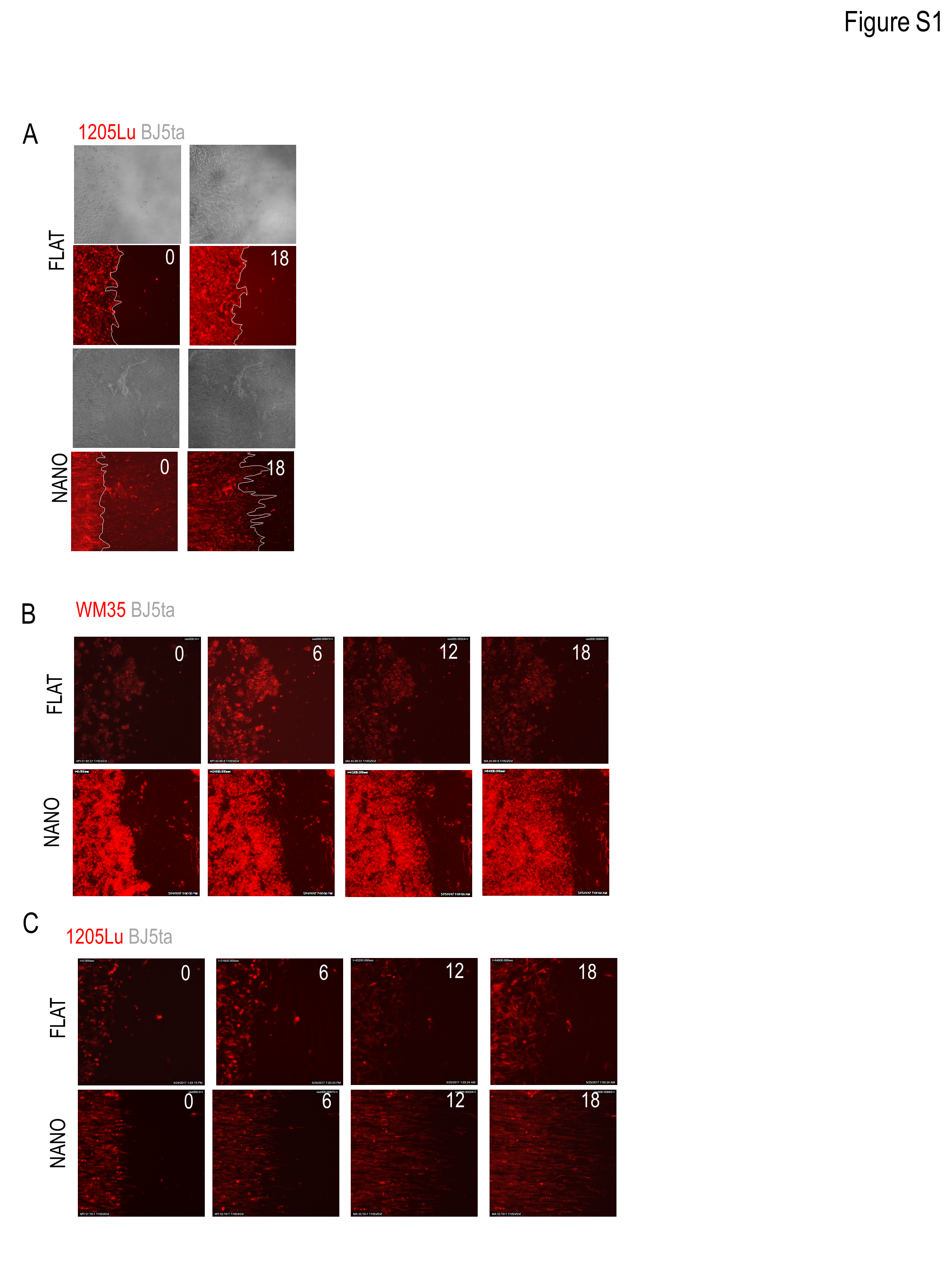

### Suppl Figure 2

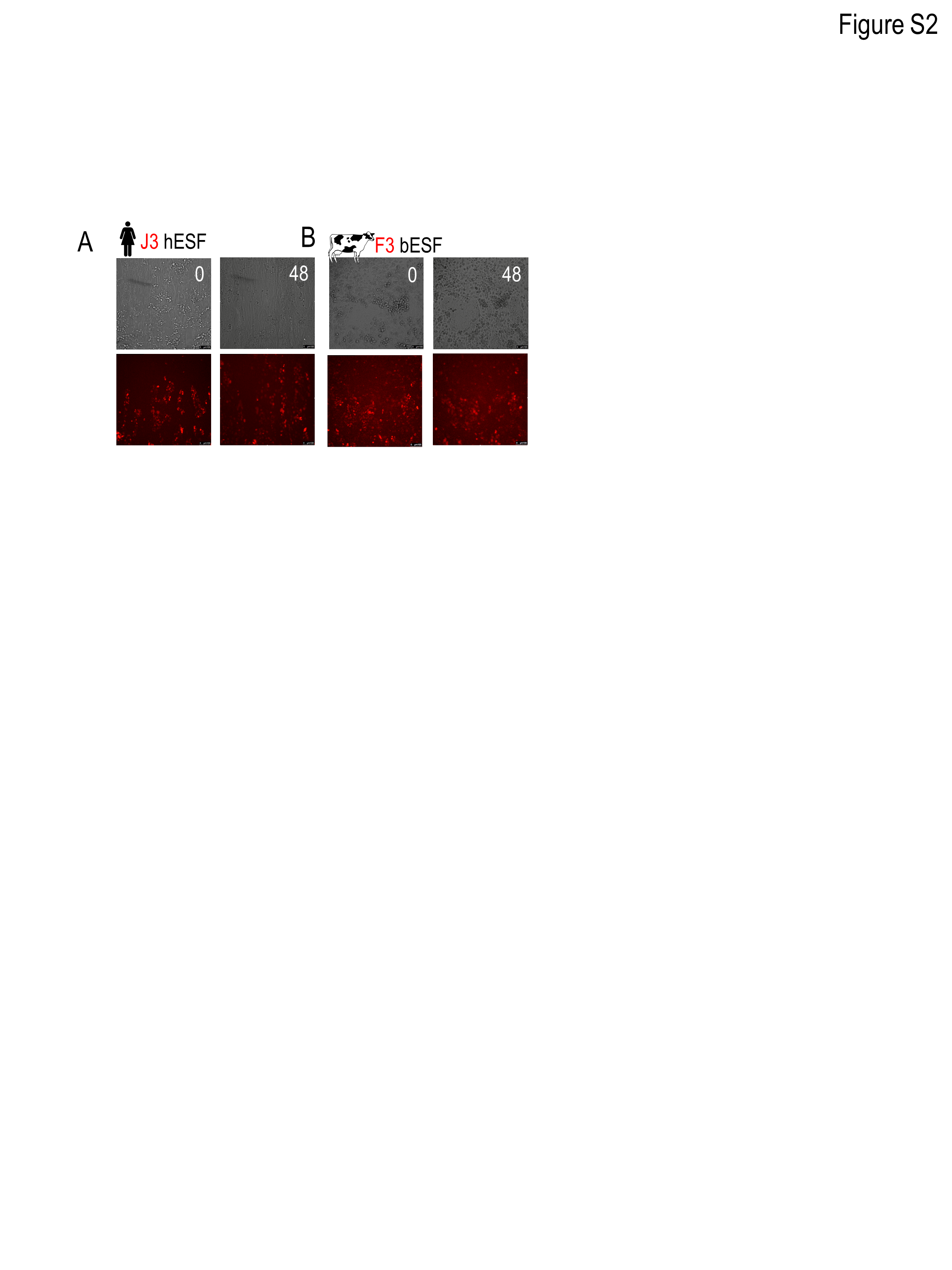

### Suppl Figure 3

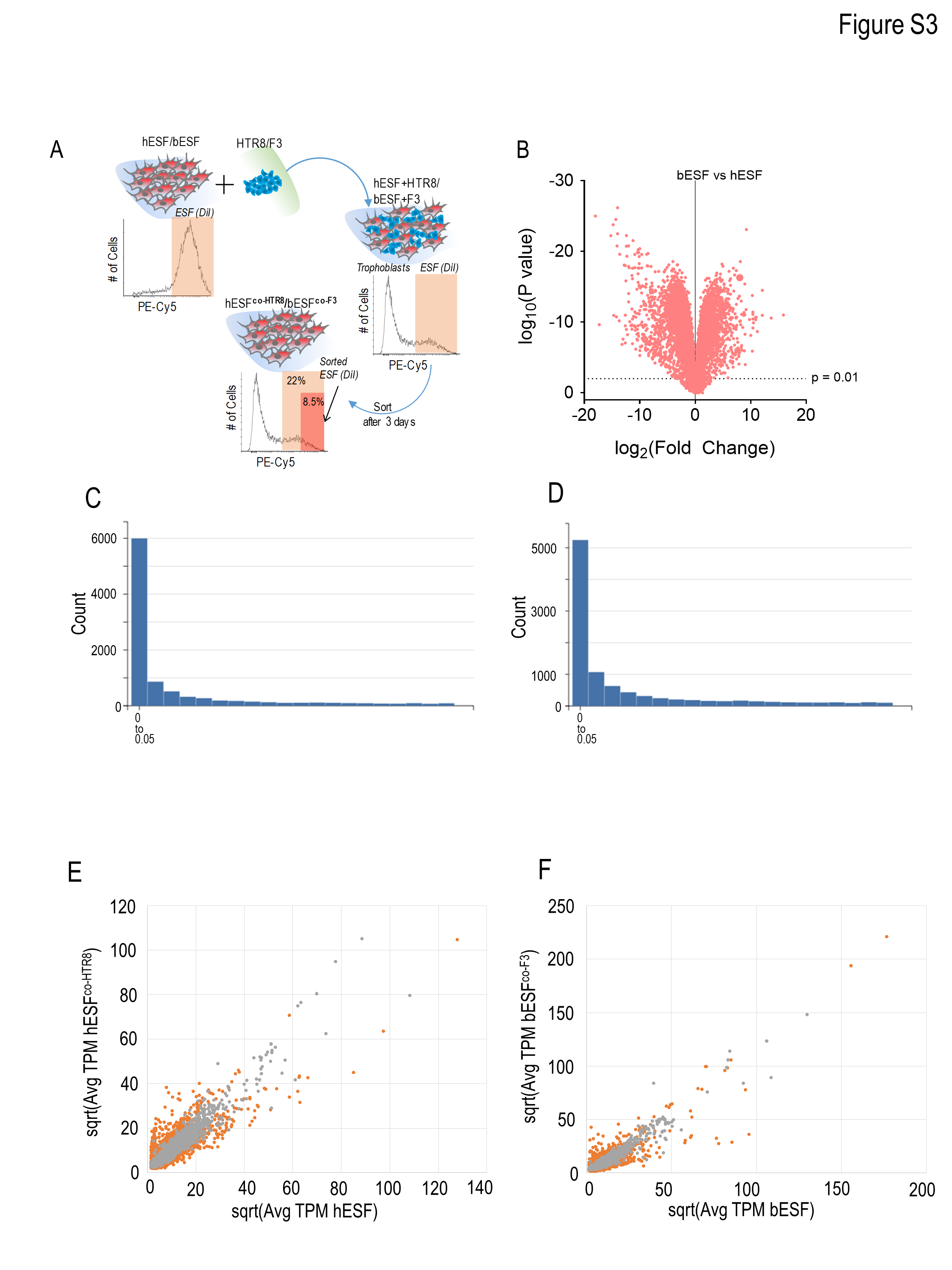

### Suppl Figure 4

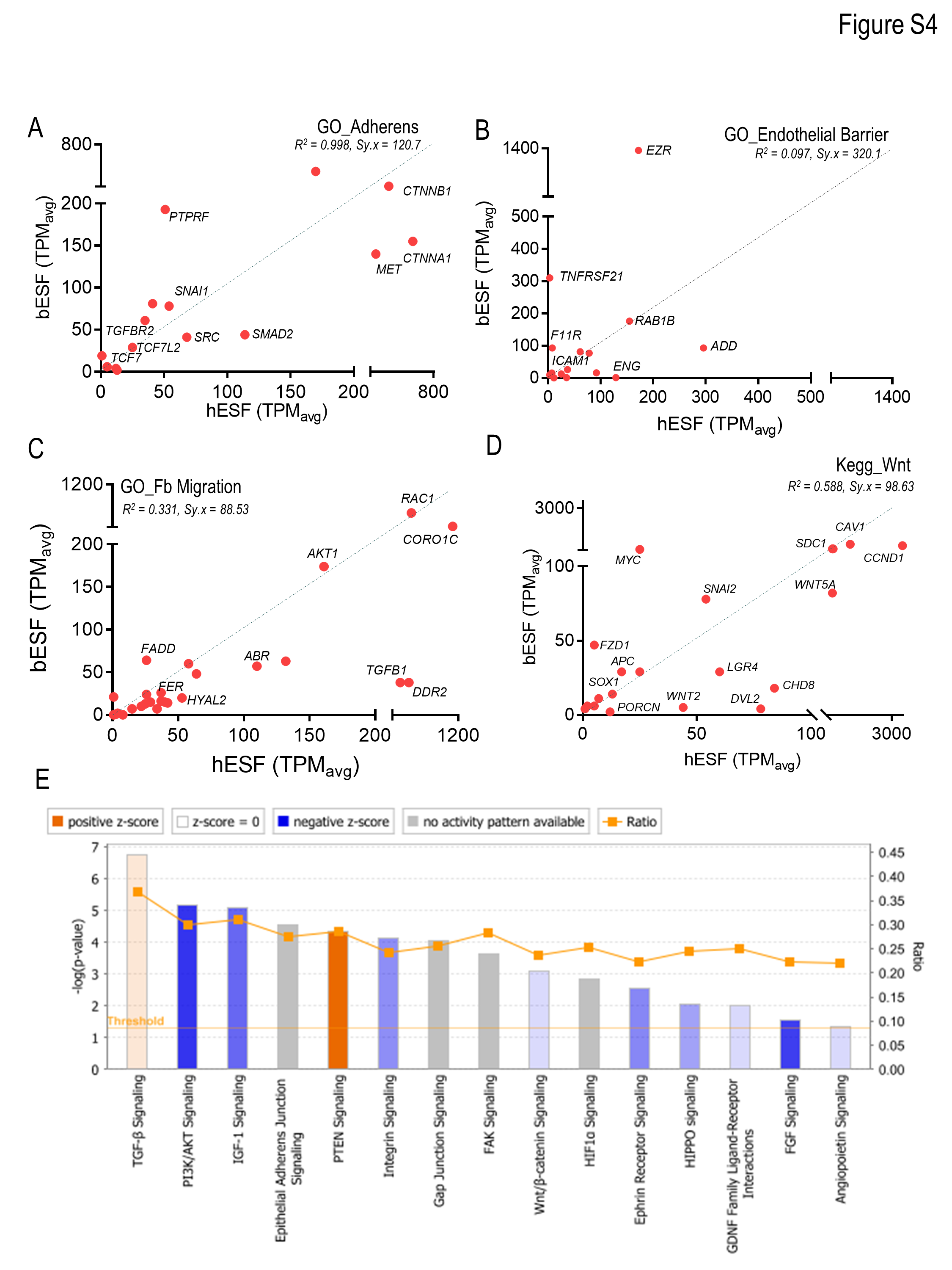

### Suppl Figure 5

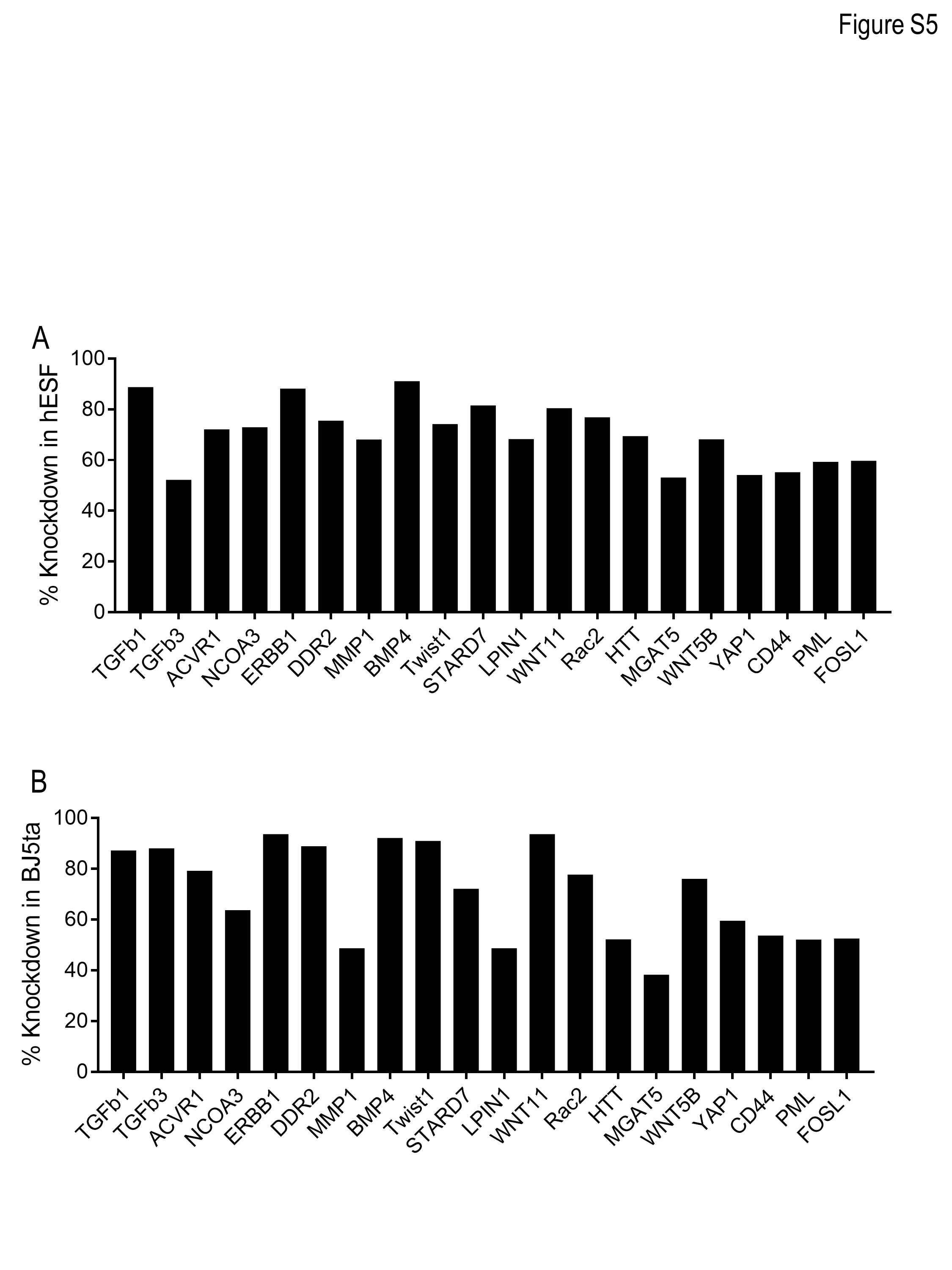
